# OSBP is a major determinant of Golgi phosphatidylinositol 4-phosphate homeostasis

**DOI:** 10.1101/2023.12.21.572879

**Authors:** Colleen P. Doyle, Liz Timple, Gerald R. V. Hammond

## Abstract

The lipid phosphatidylinositol 4-phosphate (PI4P) plays a master regulatory role at Golgi membranes, orchestrating membrane budding, non-vesicular lipid transport and membrane organization. It follows that harmonious Golgi function requires strictly maintained PI4P homeostasis. One of the most abundant PI4P effector proteins is the oxysterol binding protein (OSBP), a lipid transfer protein that exchanges trans Golgi PI4P for ER cholesterol. Although this protein consumes PI4P as part of its lipid anti-porter function, whether it actively contributes to Golgi PI4P homeostasis has been questioned. Here, we employed a series of acute and chronic genetic manipulations, together with orthogonal targeting of OSBP, to interrogate its control over Golgi PI4P abundance. Modulating OSBP levels at ER:Golgi membrane contact sites produces reciprocal changes in PI4P levels. Additionally, we observe that OSBP has a high capacity for PI4P turnover, even at orthogonal organelle membranes. However, despite also visiting the plasma membrane, endogenous OSBP makes no impact on PI4P levels in this compartment. We conclude that OSBP is a major determinant of Golgi PI4P homeostasis.

## Introduction

Oxysterol binding protein (OSBP)-related proteins (ORPs) facilitate the non-vesicular transfer of sterols and phosphatidylserine (PS) from their site of synthesis in the endoplasmic reticulum (ER) to target organelles, such as the Golgi, plasma membrane or endosomes (Subra et al., 2023; Nakatsu and Kawasaki, 2021). Since the target organelles typically maintain a higher concentration of these lipids than the ER, transport occurs against a chemical gradient, requiring an input of energy. There is accumulating evidence that this energy comes from ORPs’ anti-port of phosphatidylinositol 4-phosphate (PI4P) molecules from target organelles to the ER, down a concentration gradient (Saint-Jean et al., 2011; Mesmin et al., 2013; Chung et al., 2015; Filseck et al., 2015). PI 4-kinases maintain PI4P concentrations in the target organelle, whereas the lipid phosphatase SAC1 degrades it in the ER (Mesmin et al., 2013; Chung et al., 2015). Ultimately, the energy for sterol and PS transport comes from the hydrolysis of ATP by the PI 4-kinases. In turn, the enrichment of PS and/or sterols in target organelles can facilitate other changes in membrane composition. For example, PS is enriched at repairing lysosomes by ORPS 9, 10 and 11, which in turn recruits ATG2 to facilitate bulk phospholipid import (Tan and Finkel, 2022). This places PI4P and the ORPs at the center of non-vesicular lipid transport.

On the other hand, PI4P also serves more canonical phosphoinositide signaling roles, facilitating recruitment and/or activation of peripheral membrane proteins. Examples include clathrin adapters at the trans Golgi (Wang et al., 2003; Szentpetery et al., 2010), structural proteins that integrate the cytoskeleton with the Golgi (Dippold et al., 2009) and endosomal protein PLEKHM2 (Levin-Konigsberg et al., 2019). Notably, many ORPs themselves contain a PI4P-binding pleckstrin homology (PH) domain through which they attach to their target organelles. Therefore, maintenance of PI4P concentrations in organelle membranes, particularly the Golgi, is essential beyond roles in lipid anti-port. This poses the question: to what extent is ORP-mediated PI4P anti-port a major source of PI4P turnover, and does it play a role in organelle PI4P homeostasis?

Quantitative proteomics has recently shown that OSBP is the most abundant ORP in the HEK293 model cell line, with approximately 210,000 copies per cell (Cho et al., 2022). Experiments that acutely inhibited OSBP with a small molecule inhibitor in retinal pigment epithelial (RPE-1) cells suggested consumption of approximately 80% of the Golgi PI4P pool by this protein (Mesmin et al., 2017). Nevertheless, cells possess other mechanisms for PI4P degradation. SAC1 traffics between the ER and Golgi membranes, and its traffic to the Golgi in response to serum starvation has been shown to dramatically decrease Golgi PI4P levels and blunt secretion (Blagoveshchenskaya et al., 2008). Endosomal PI4P is hydrolyzed by the peripheral membrane phosphatase, SAC2 (Nakatsu et al., 2015; Hsu et al., 2015), and plasma membrane Synaptojanins 1 and 2 possess SAC domains that can hydrolyze PI4P (Chung et al., 1997; Nemoto et al., 2001). Furthermore, studies in HeLa and CHO cells reported no effect of OSBP knock-down on PI4P levels (Venditti et al., 2019; Goto et al., 2016). Recently, experiments in HeLa cells have shown ORPs 9 and 10 are essential for regulation of PI4P levels, with their loss leading to hyper-recruitment and activity of OSBP (Venditti et al., 2019; Naito et al., 2023). Recently, experiments in HeLa cells have shown ORPs 9 and 10 are essential for regulation of PI4P levels, rather than OSBP (Venditti et al., 2019; Naito et al., 2023). ORP9/10 loss then leads to hyper-recruitment and activity of OSBP.

Collectively, whereas OSBP is crucial for traffic of cholesterol to the Golgi, its role in Golgi PI4P turnover is less clear. Is PI4P anti-port by OSBP a major route of PI4P turnover? In other words, does OSBP play a major role in Golgi PI4P homeostasis? We have addressed this question in the current study. We employed an array of experiments to acutely or chronically deplete OSBP activity in HEK293A cells, as well as targeting the protein to orthogonal compartments. We could then assay PI4P homeostasis in living cells as a read out. We demonstrate that OSBP has a large capacity to facilitate PI4P turnover and is a major player in PI4P homeostasis. However, it makes little contribution to PI4P turnover in another organelle that it visits: the plasma membrane.

## Results

### Manipulation of OSBP activity reveals a contribution to PI4P homeostasis

We measured PI4P levels in intact, living cells using P4Mx1 (Hammond et al., 2014), since this biosensor does not perturb Golgi morphology, is exquisitely specific for PI4P in all cellular membranes, yet is relatively low affinity and therefore sensitive to both increased and decreased PI4P levels. To acutely test for a function for OSBP in Golgi PI4P turnover, we turned to the potent inhibitor of OSBP lipid transfer activity, OSW-1 (Burgett et al., 2011; Mesmin et al., 2017). This compound blocks cholesterol and PI4P binding by the OSBP-related domain (ORD). Consequently, this should cause PI4P accumulation in the Golgi if PI4P exchange by OSBP is a major route of PI4P catabolism (**Fig. 1A**). Exactly as reported previously (Mesmin et al., 2017), we found that inhibition of OSBP led to a rapid increase in Golgi PI4P levels (**Fig. 1B**), completing within 20 minutes (**Fig. 1C**). The increase was highly significant by area-under-the-curve analysis (**Fig. 1D**).

**Figure 1:**
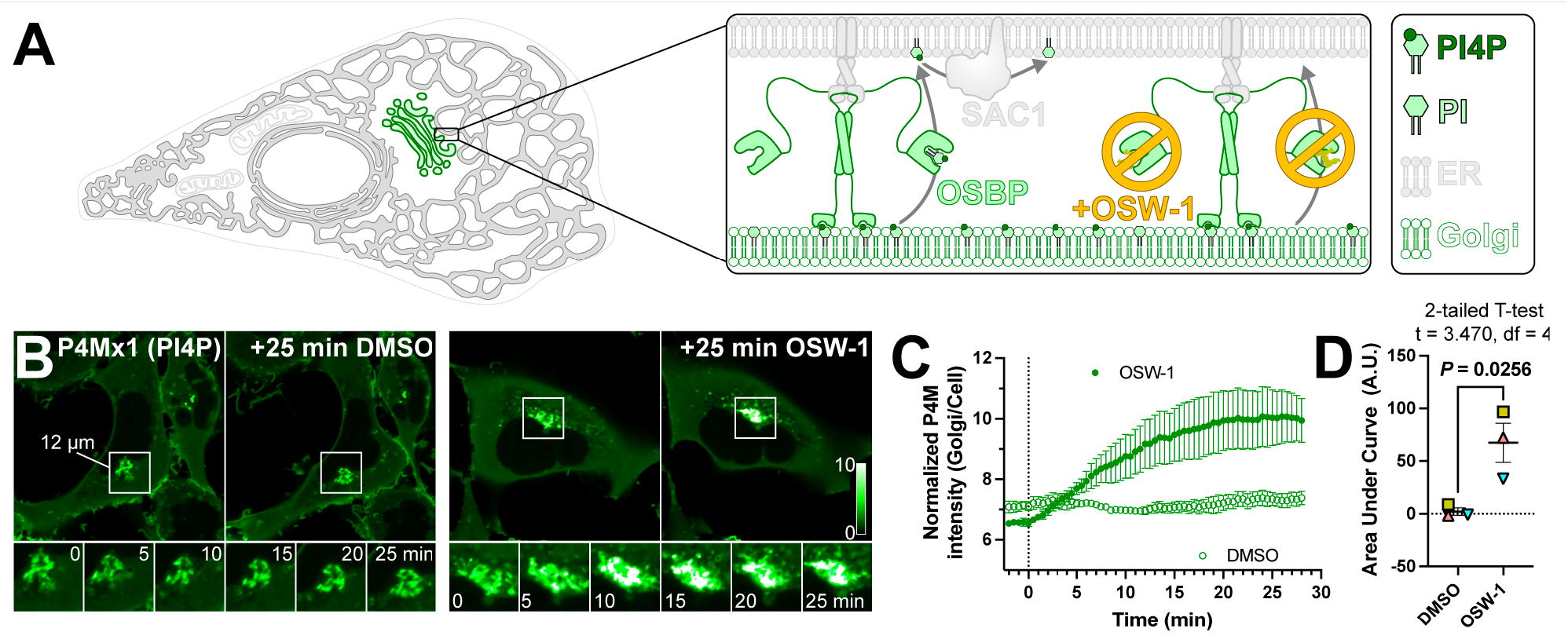
Acute pharmacologic inhibition of OSBP increases Golgi PI4P levels. (**A**) OSBP transfers PI4P molecules from Golgi membranes to the ER for degradation at membrane contact sites, which can be inhibited by OSW-1. (**B**) HEK293A cells expressing EGFP-P4Mx1 PI4P biosensor were imaged by confocal time-lapse microscopy during treatment with either 20 nM OSW-1 or 0.02% DMSO as vehicle control. Insets at bottom show intermediate time points of the boxed region. The intensity scale is normalized to the pixel average for each cell. (**C**) quantification of the experiment shown in B; data are grand means ± s.e.m. of 3 experiments, measuring 10 cells each. (**D**) Analysis of the area under the curve for the data presented in C; matching symbols represent the area for each experiment. Means ± s.e.m. are also indicated.

If acute inhibition of OSBP leads to rapid accumulation of Golgi PI4P, why have prior reports not shown similar accumulations after siRNA-mediated knock-down of OSBP (Goto et al., 2016; Venditti et al., 2019)? We reasoned that the difference could be timing: OSW-1 acutely inhibits the OSBP protein within seconds, leading to a response within minutes (**Fig. 1**). RNAi, on the other hand, leads to depletion of the protein over 2-3 days, perhaps allowing the cell to adapt PI4P homeostasis back to the steady state via other mechanisms. We therefore wanted to devise an experiment whereby OSBP could be acutely removed from the Golgi. To this end, we first employed RNAi to deplete endogenous OSBP, and made an RNAi-resistant human OSBP fusion protein to replace it, linked to mCherry and the FKBP12 protein (**Fig. 2A**). We found the FKBP-OSBP protein itself did not affect Golgi PI4P, but in contrast to the prior reports in HeLa and CHO cells, OSBP siRNA did lead to detectable accumulation of Golgi PI4P in HEK293A cells (**Figs. 2B, C**). Reassuringly, this was completely reversed by expression of the RNAi-resistant FKBP-OSBP (**Figs. 2B,C**).

**Figure 2:**
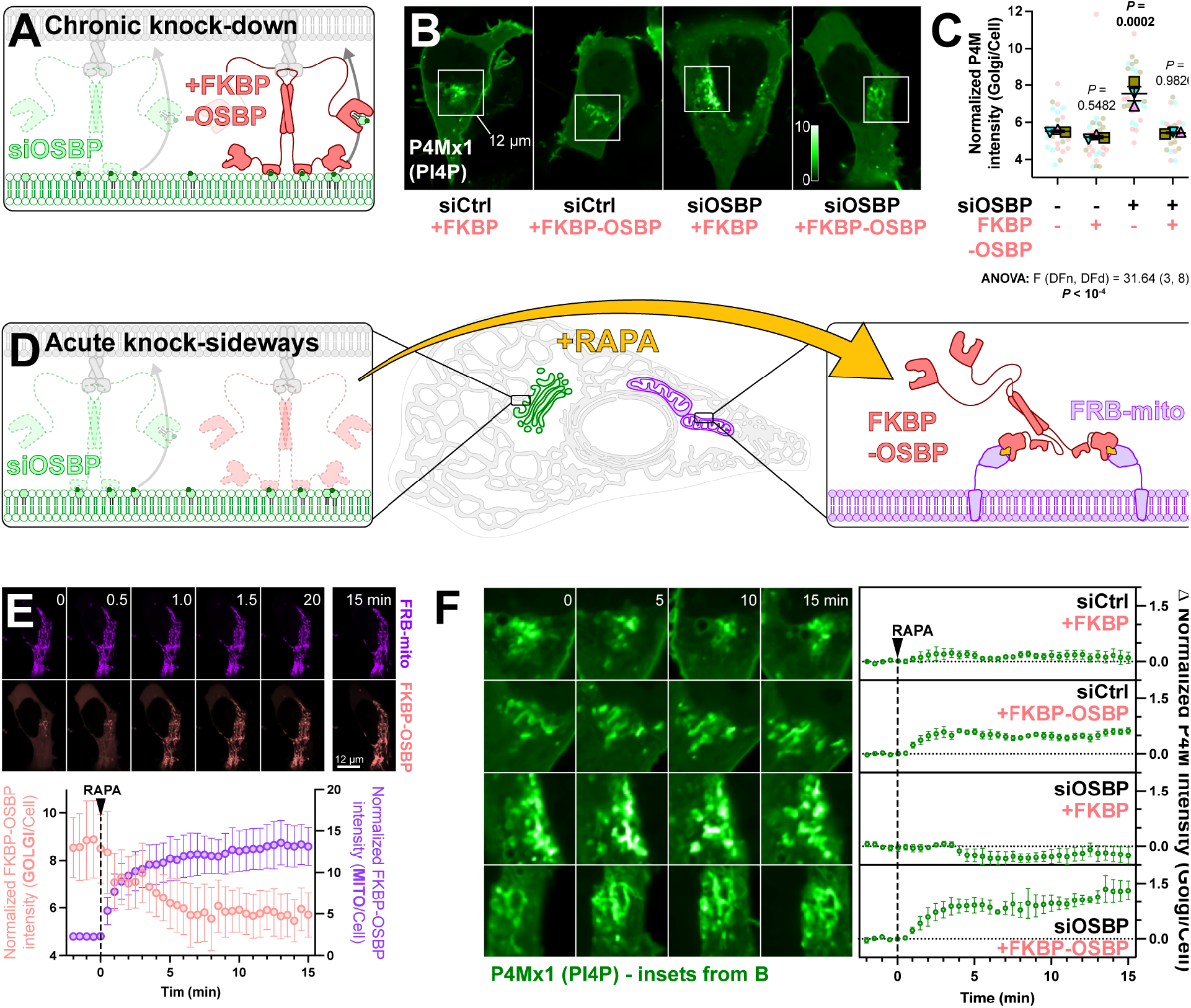
Chronic and acute depletion of OSBP increases Golgi PI4P levels. (**A**) Experimental strategy using knock-down of endogenous OSBP with siRNA and rescue with FKBP-OSBP expression. (**B**) Confocal sections of representative cells expressing PI4P biosensor P4Mx1 after co-transfecton with the indicated siRNA and mCherry-tagged protein. (**C**) super plot showing the quantification of Golgi fluorescence normalized to whole confocal section of each cell under the indicated transfection conditions. Pale symbols represent individual cell measurements (5-10 per experiment), whereas larger symbols represent means of each of 3 experiments. Both symbol types are color matched by experiment. Grand mean ± s.e. is also indicated. Results of a one-way ANOVA are indicated; *P* values above the data are from Dunnett’s multiple comparisons test relative to siCtrl + FKBP. (**D**) Experimental strategy using acute sequestering of FKBP-OSBP to mitochondria by rapamycin-induced dimerization with FRB-mito (“knock-sideways”) in cells wherein endogenous OSBP was depleted by siRNA. (**E**) Rapid sequestering of mCherry-FKBP-OSBP to mitochondria. Images show confocal time-lapse of a cell expressing FKBP-OSBP and FRB-mito. Quantification below shows grand means ± s.e. of 3 experiments. (**F**) Effect of acute OSBP knock-sideways on PI4P biosensor accumulation at the Golgi. Images show time-lapse of the inset region from panel B for the indicated times after knock-sideways initiation with 1 µM rapamycin. The graphs show the change in Golgi fluorescence normalized to whole confocal section; data are grand means ± s.e. of 3 independent experiments.

Having shown that our FKBP-OSBP was functional, we paired the system with a mitochondrially-targeted fragment of mTOR that binds rapamycin (FRB). This enables rapamycin-induced dimerization of FKBP-OSBP with mitochondrially-restricted FRB, and hence rapid “knock-sideways” of the OSBP (Robinson et al., 2010). Functionally, this should equate to a rapid depletion of the protein from the Golgi within minutes (**Fig. 2D**). Indeed, rapamycin addition led to the rapid recruitment of cytosolic and Golgi FKBP-OSBP to the mitochondria, and a concomitant depletion of Golgi OSBP (**Fig. 2E**). This completed within 5 mins – much faster than the 72 hours over which RNAi was conducted. Knock-sideways of the FKBP-only control had no effect on Golgi PI4P pools in non-targeting or OSBP knock-down cells, but we saw a rapid (within 4 minutes) increase in Golgi PI4P when FKBP-OSBP was knocked sideways in OSBP knock-down cells (**Fig. 2F**, bottom panel). We saw a similar but more muted effect in cells treated with control RNAi (**Fig. 2F**, second from top). We postulate that this occurs due to dimerization of OSBP (Mora et al., 2021), leading to knock-sideways of some endogenous OSBP in these cells when dimerized with FKBP-OSBP.

Although we could see increased Golgi PI4P in response to rapid (minutes) or slow (days) removal of OSBP activity, we did not observe the converse: depleted PI4P after OSBP over-expression (**Fig. 2B**). We reasoned this is because endogenous VAP-A/B ER receptors are saturated, therefore unable to target additional OSBP to ER:Golgi contact sites for productive PI4P transfer (**Fig. 3A**). We therefore hypothesized that combined over-expression of OSBP and VAP-B would deplete Golgi PI4P. We compared over-expression of these proteins to Sec61β and the GRIP domain from Golgin A4, as controls for transmembrane ER and peripheral trans Golgi network (TGN) proteins, respectively. As shown in **Figs. 3B and C**, only combined over-expression of VAP-B and OSBP proteins led to substantial depletion of Golgi PI4P. Neither protein alone led to PI4P depletion (**Fig. 3B, C**).

**Figure 3:**
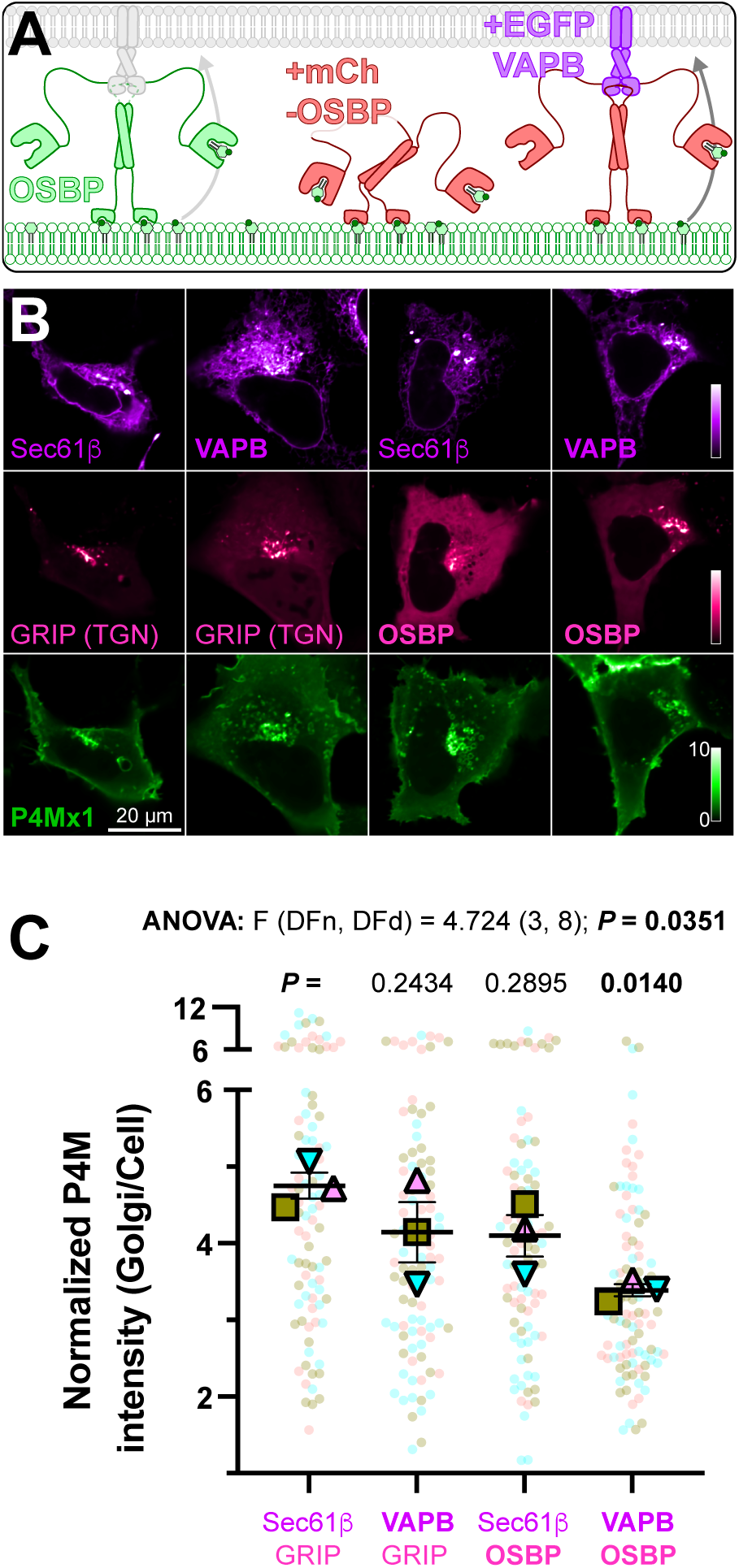
OSBP over-expression decreases Golgi PI4P when VAPB is co-expressed. (**A**) Over-expressed OSBP could be unable to further deplete Golgi PI4P since there are no available endogenous VAP proteins unless these are also over-expressed. (**B**) Confocal sections of HEK293A cells over-expressing iRFP713-P4Mx1 PI4P biosensor, EGFP-VAPB (or AcGFP-Sec61β as control) and mCherry-OSBP (or the GRIP domain of Golgin A4 as control). (**C**) super plot showing the quantification of Golgi fluorescence normalized to whole confocal section of each cell under the conditions detailed in B. Pale symbols represent individual cell measurements (27-32 per experiment), whereas larger symbols represent means of each of 3 experiments. Both symbol types are color matched by experiment. Grand mean ± s.e. is also indicated. Results of a one-way ANOVA are indicated; *P* values above the data are from Dunnett’s multiple comparisons test relative to the Sec61β + GRIP control.

### Endogenous tagging of OSBP reveals turnover of ectopic PI4P pools

To further test for a function of PI4P transfer by OSBP in homeostasis of the lipid, we designed a strategy to orthogonally target endogenous OSBP in living cells, and test its effect on PI4P levels in that orthogonal compartment. As a first step, we needed an approach to monitor endogenous OSBP in living cells. To this end, we employed genomic tagging of OSBP using a split-neonGreen tag (NG2, **Fig. 4A**), as recently demonstrated by the OpenCell project (Cho et al., 2022). Using the guide RNA and homology repair template described by OpenCell, we generated HEK293A cells with neonGreen fluorescence decorating juxta-nuclear structures, consistent with the trans-Golgi network in these cells (**Fig. 4B**), exactly as reported (Cho et al., 2022). To confirm OSBP was the tagged protein, we treated these cells with the siRNA pool against OSBP and found green fluorescence was depleted (**Fig. 4B, C**). Additionally, we could show that the TGN-like staining of NG2-OSBP cells co-localized with immunostaining against OSBP in fixed cells (**Fig. 4D**).

**Figure 4:**
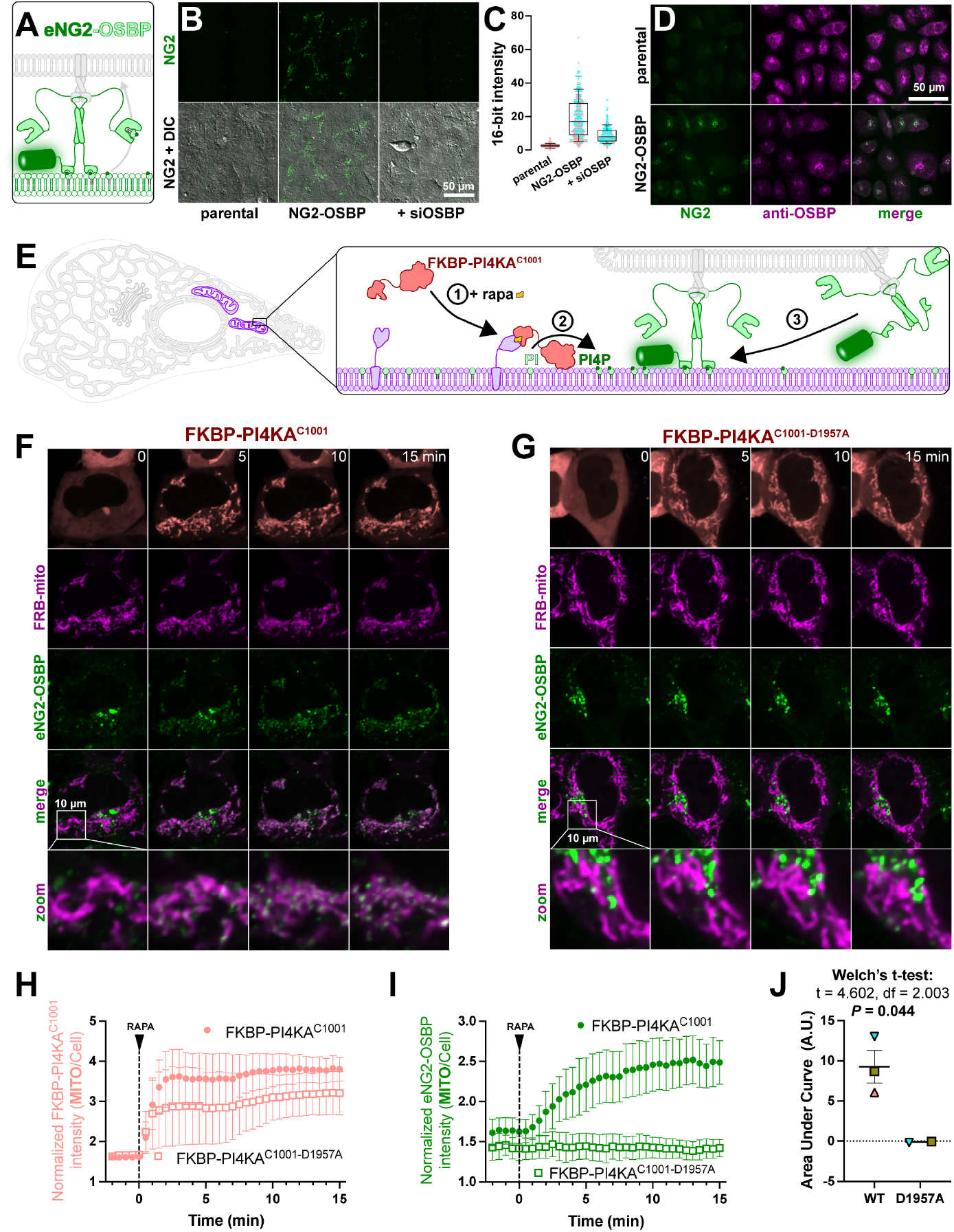
Translocation of endogenous OSBP to mitochondrial outer membranes in response to ectopic PI4P synthesis. (**A**) Genomic tagging of OSBP with a split NeonGreen2 tag. (**B**) Representative confocal micrographs of parental HEK293A cells, NG2-OSBP knock in cells and NG2-OSBP knock in cells transfected for 48h with short interfering RNA against OSBP. DIC images were subject to a Fourier bandpass filter to correct for non-uniform illumination. NG2 channels are shown at identical excitation and gain settings. (**C**) Average intensity of cells like those shown in B from two experiments (measuring between 147 and 198 cells per condition from random fields), showing depletion of NG2 signal in NG2-OSBP cells by siOSBP. (**D**) Co-localization of NG2-OSBP signal with antibody staining against endogenous OSBP. Fields are representative of two experiments. (**E**) Experimental strategy using chemically induced dimerization to recruit a PI4K to the mitochondrial outer membrane to synthesize PI4P, and the hypothesized recruitment of OSBP via its PI4P-binding PH domains. (**F**) Confocal time-lapse of a representative NG2-OSBP cells after mitochondrial PI4P synthesis is induced with rapamycin. Note the appearance of green puncta on the mitochondria marker by FRB-mito (magenta). (**G**) as F, except the catalytically inactive PI4KA mutant, D1957A is used and no green puncta appear at mitochondria. (**H-I**) Quantification of the fluorescence intensity of FKBP-PI4KAC1001 (H) and NG2-OSBP (I) at mitochondria after rapamycin induced dimerization of FKBP with FRB-mito. Data are grand means ± s.e. of two (PI4KA) or three (D1957A) independent experiments quantifying 7-9 cells each. (**J**) Area under the curve calculation for the data presented in I showing significant recruitment of NG2-OSBP by PI4KA (WT) but not the D1957 mutant.

Next, we turned to a method to orthogonally target OSBP in cells. OSBP’s localization to the Golgi depends on an interaction of its N-terminal pleckstrin homology domain with PI4P (Levine and Munro, 2002; Balla et al., 2005). Several studies have recently shown that targeting of an FKBP-tagged PI 4-kinase to the mitochondrial outer membrane leads to ectopic accumulation of PI4P in this organelle (Doyle et al., 2023; Pemberton et al., 2020; Zewe et al., 2020). We therefore reasoned that this mitochondrial PI4P might recruit endogenous OSBP (**Fig. 4E**). Indeed, we found that recruitment of an FKBP-tagged c-terminal fragment of PI4KA (Zewe et al., 2020) to mitochondria caused the redistribution of NG2-OSBP from the juxta-nuclear region to mitochondria, which became decorated with NG2-OSBP puncta (**Fig. 4F**). These are likely to be sites of ER-mitochondria contact. In contrast, recruitment of OSBP was not apparent when a catalytically inactive PI4KA was used (**Fig. 4G**), despite similarly robust recruitment of the kinase (**Fig. 4H**). Indeed, quantitative analysis of the fluorescence intensity at mitochondria showed no change with the inactive mutant, compared to rapid (within 5 min) recruitment of NG2-OSBP using active PI4KA (**Fig. 4I**). Area-under-the-curve analysis showed substantial and significant recruitment of NG2-OSBP to mitochondria by PI4KA compared to the inactive mutant (**Fig. 4J**).

Since OSBP could be orthogonally targeted to mitochondria by PI4P, we next wanted to ask the question: does OSBP then play a role in turnover of this ectopic PI4P pool? To address this question, we employed two approaches (**Fig. 5A**). Firstly, we tested if ectopic PI4P accumulaton at mitochondria was impacted by OSBP inhibition (using OSW-1). Secondly, we tested whether PI4P turnover requires OSBP, as it does at the Golgi. For this latter test, we turned to a reversible system for synthesis of ectopic PI4P. Preliminary experiments used the potent and selective PI4KA inhibitor, GSK-A1 (Bojjireddy et al., 2014) to inactivate mitochondrially targeted FKBP-PI4KA. However, this approach yielded inconclusive results, because so much EGFP-P4Mx1 biosensor was released from the plasma membrane after PI4KA inhibition. Instead, we turned to a reversible chemically-induced dimerization system (Schifferer et al., 2015; Feng et al., 2014). This uses small molecule-induced dimerization between a SNAP tag and FKBP tag with the synthetic small molecule, rCD1 – in our context, recruiting FKBP-PI4KA to mitochondrially targeted SNAP (see step 1 in **Fig. 5A**). Unlike the high affinity FKBP-FRB-rapamycin ternary complex, the ternary complex of rCD1 and SNAP with FKBP has much lower affinity, such that it is readily competed by FK506, causing rapid release of FKBP-PI4KA from the SNAP-rCD1 complex (step 2a in **Fig. 5A**).

**Figure 5:**
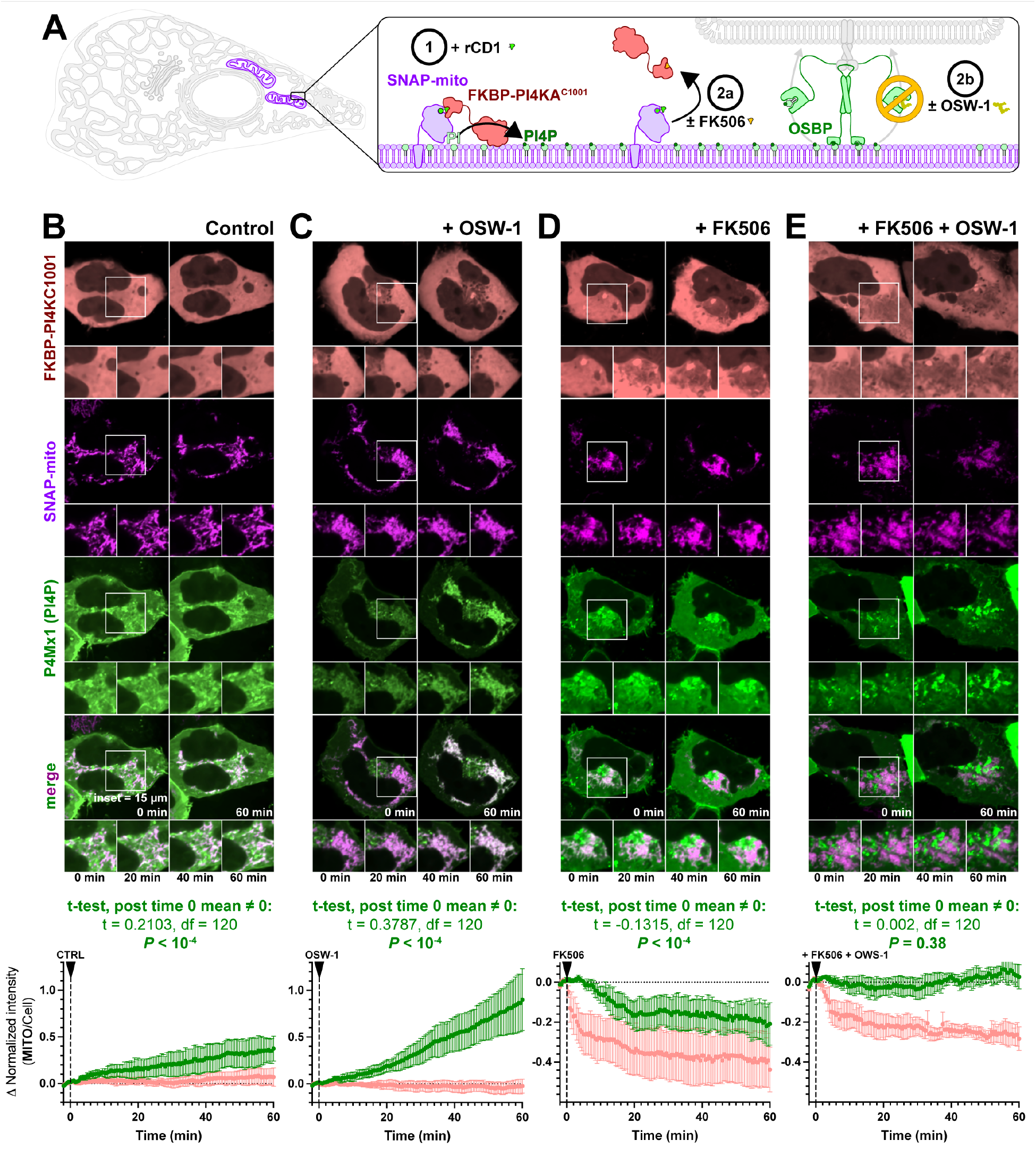
Endogenous OSBP removes ectopic PI4P from the mitochondrial outer membrane. (**A**) Cells express a mitochondrial outer membrane targeted SNAP tag, which undergoes chemically-induced dimerization with FKBP-tagged PI4KAC1001 in the presence (1 µM) of the reversible chemical dimerizer (rCD1). At the onset of the experiment, cells are imaged in the presence of either DMSO control or 1 µM FK-506 to compete the rCD1:SNAP complex from the PI4K, and thereby induce dissociation. Where indicated, cells were also treated with 20 nM OSW-1 to inhibit OSBP PI4P transfer. (B-E) Confocal images of representative cells taken during time-lapse imaging of the cells during the experiment, along with quantification of the change in mitochondrial fluorescence intensity (normalized to the whole cell) for the SNAP-PI4K_C1001_ (pink) and P4Mx1 PI4P biosensor (green). Data are grand means ± s.e. for 3 experiments analyzing 4-8 cells each. The t-test reports the probability that the post-treatment deviation of the data is not 0.

Preliminary experiments with these constructs yielded relatively inefficient recruitment of FKBP-PI4KA to mitochondria over 30 mins. Therefore, we conducted the experiment to pre-incubate cells with rCD1 for 60 mins prior to imaging, selecting cells with evidence of FKBP-PI4KA recruitment and P4Mx1 labelling at mitochondria. Control experiments in which no FK506 or OSW-1 were added revealed a continued, slow increase in mitochondrial PI4P over the subsequent 60 mins of the experiment (**Fig. 5B**). Inhibition of OSBP activity using OSW-1 caused a much more dramatic increase in PI4P over this time frame, suggesting OSBP is normally opposing PI4P accumulation on the mitochondria (**Fig. 5C**). On the other hand, addition of FK506 to cells led to rapid (within 2 mins) loss of FKBP-PI4KA, followed by a somewhat lagging decline in PI4P levels over the following 60 mins (**Fig. 5D**). Co-treatment of cells with OSW-1 to block OSBP activity at the same time as FK506 addition completely blocked the PI4P decline, despite robust reduction in FKBP-PI4KA (**Fig. 5E**). Therefore, it appears that OSBP activity is required to deplete PI4P from mitochondria after removal of PI4K activity. The most parsimonious explanation for these results is that OSBP is able to transfer PI4P from the mitochondrial outer membrane back to the ER for degradation by SAC1. The data underscore a strong capacity of endogenous OSBP to contribute to PI4P turnover in cells.

### OSBP does not measurably contribute to plasma membrane PI4P turnover

If OSBP impacts turnover of PI4P in the Golgi and is capable of turning it over in an orthogonal compartment after artificial induction of PI4P synthesis, we next asked if OSBP/ was able to contribute to turnover of PI4P in other PI4P-rich membranes? The plasma membrane (PM) contains the largest pool of PI4P in the cell (Hammond and Balla, 2015). We therefore tested for a potential role for OSBP in impacting PI4P turnover here (**Fig. 6A**). Imaging the PM of NG2-OSBP cells using total internal reflection fluorescence microscopy (TIRFM) revealed the presence of uniform neonGreen puncta associated with the PM (**Fig. 6B**). Time lapse imaging showed that these puncta stayed associated with the PM for a few seconds (**Fig. 6B**, kymograph). Analysis of the puncta full-width at half maximum (FWHM) showed they were diffraction-limited in size, and the intensity was a lognormal distribution, typically emitting a few hundred photons in the 50 ms exposure period (graphs in **Fig. 6B**). This is consistent with the characteristics of single mNeonGreen protein molecules (Shaner et al., 2013), though we did not calibrate the measurement to distinguish monomers from dimers, which would be expected for OSBP. Co-localization with markers of the ER (Sec61β) or ER-PM contacts sites using MAPPER (Chang et al., 2013) showed that OSBP puncta were highly enriched at these structures when viewed in TIRFM (**Fig. 6C**), consistent with association between OSBP and ER-localized VAP proteins. As a negative control, no enrichment of OSBP was seen with clathrin labelled structures, imaged using mCherry-tagged clathrin light chain (**Fig. 6C**). Puncta were present at relatively low density, averaging 19.7 ± 1.8 puncta/100 µm2 (mean ± s.e., n = 24 cells) of membrane in clathrin light chain co-expressing controls.

**Figure 6:**
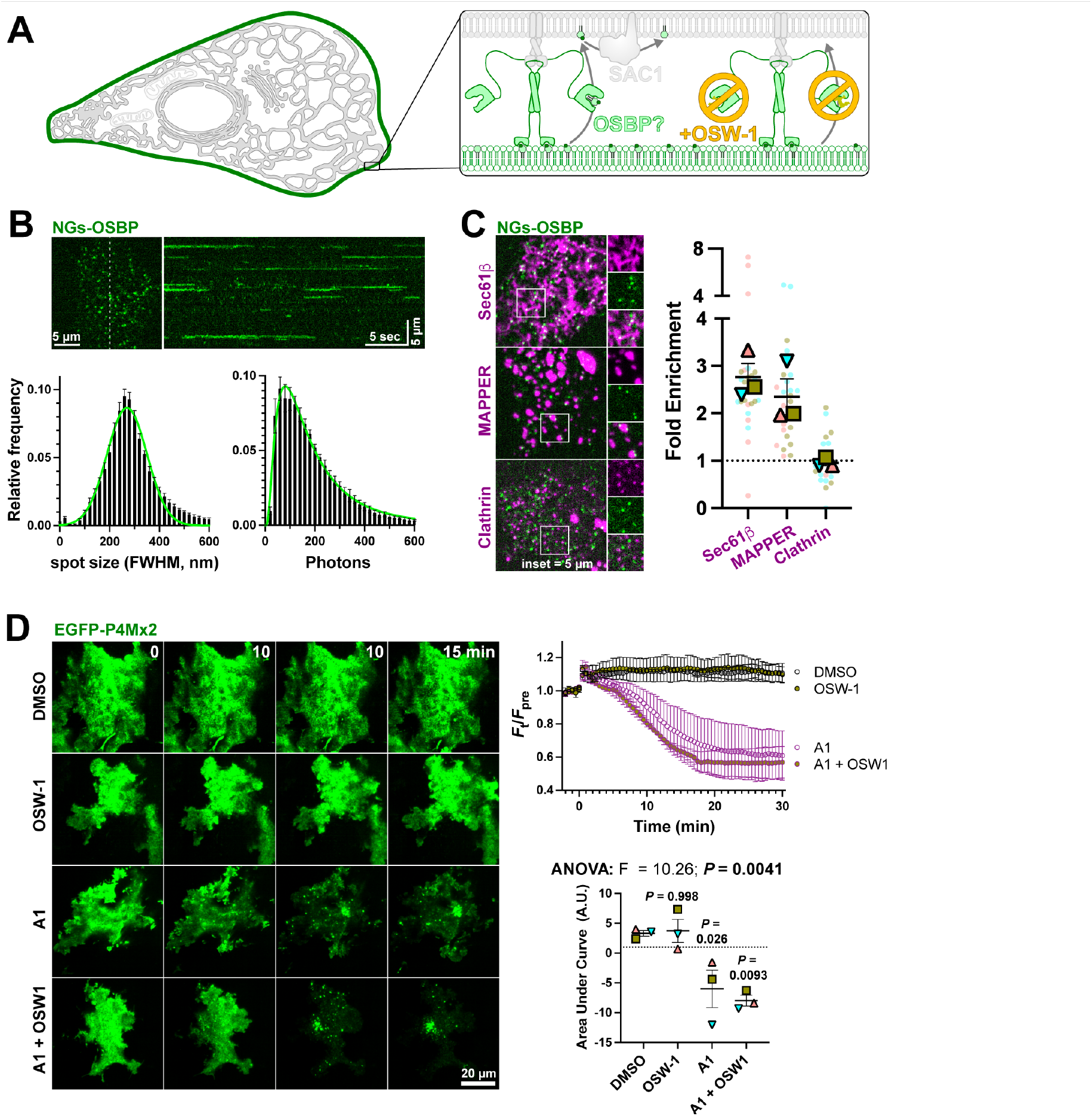
Endogenous OSBP visits the PM at ER contact sites, but makes no measurable impact on PM PI4P turnover. (**A**) Hypothesized recruitment of OSBP to ER:PM contact sites through PM PI4P, and its inhibition by OSW-1. (**B**) TIRFM of NG2-OSBP cells shows numerous puncta at the PM; the kymograph at right shows the view along the dashed line over 30 sec, showing puncta persist for several seconds. The graphs show the puncta full-width at half maximum and intensity (in photons), exhibiting values consistent with single molecules. Data are means ± 95% C.I. from 9 cells from a representative experiment. Trend lines fit a normal or log-normal distribution. (**C**) NG2-OSBP particles viewed in TIRFM visit the PM mainly at ER:PM contact sites. Images show co-localization of NG2-OSBP with mCherry-Sec61β (ER marker), -MAPPER (ER:PM contact site marker) or -clathrin light chain (negative control). The graphs shows the relative enrichment of puncta at these markers; pale symbols represent individual cell measurements (3-10 per experiment), whereas larger symbols represent means of each of 3 experiments. Both symbol types are color matched by experiment. Grand mean ± s.e. is also indicated. (**D**) OSBP does not measurably impact PM PI4P turnover after inhibiting the PM PI4K with A1. COS-7 cells expressing the EGFP-P4Mx2 PI4P biosensor were imaged in TIRFM and treated with the indicated compounds. The time course plot shows TIRFM fluorescence over time, and is the grand mean ± s.e. of three independent experiments (10-13 cells per experiment). Area under the curve calculation is shown for these data. Individual points show means of each experiment; grand mean ± s.e. is also shown. Results of a one-way ANOVA are indicated; *P* values above the data are from Sidak’s multiple comparisons test relative to the DMSO control.

Since OSBP labels the PM at sites of ER:PM contact (defined by the presence of MAPPER), we next asked if inhibition of OSBP would slow the turnover of PM PI4P? PI4P turnover becomes apparent once the major PM PI4K, PI4KA, is inhibited with GSK-A1 (**Fig. 6D**). However, no slowing of PI4P decline was seen with the combined addition of GSK-A1 and OSW-1, nor was enhanced PM PI4P observed with OSW-1 addition alone (**Fig. 6D**). Therefore, despite detectable visits by endogenous OSBP to the PM, no measurable effects on PM PI4P turnover by this protein could be detected. This is in contrast to known effects of other OSBP family members ORP2, 5 and 8 (Sohn et al., 2018; Doyle et al., 2023).

## Discussion

We designed these experiments to answer the question: Is PI4P anti-port by OSBP a major route of PI4P turnover? Collectively, our result strongly indicate the answer is yes. Chronic knockdown and acute knock-sideways of OSBP leads to accumulation of Golgi PI4P, as does a small molecule inhibitor of OSBP, OSW-1 (**Figs. 1 & 2**). Conversely, over-expression of OSBP can decrease Golgi PI4P, so long as its ER receptor, VAP-B, is co-expressed in these cells (**Fig. 3**). Orthogonal targeting of endogenous OSBP to the mitochondrial outer membrane reveals this protein is sufficient to mediate turnover of an ectopically generated PI4P pool (**Figs. 4 & 5**). On the other hand, despite visiting the plasma membrane, endogenous OSBP does not make a measurable impact on PI4P in this organelle (**Fig. 6**).

Can these data be reconciled with prior reports arguing that OSBP is not a major player in PI4P turnover? Firstly, we consider direct PI4P turnover by Golgi localized SAC1. Although SAC1 traffics to the Golgi, it does so predominantly under starvation conditions during which biosynthetic secretory traffic is halted (Blagoveshchenskaya et al., 2008). Presumably, this would also eliminate Golgi targeting of OSBP and ORPs 9-11 via PI4P binding to their PH domains, shutting down lipid anti-port. This would make sense in a cell that is already shutting down secretion, since the fate of most PS and cholesterol is to traffic onwards through the secretory pathway. Next, what about alternative ORPs being responsible for PI4P homeostasis? Intriguingly, whereas OSBP is the most abundant ORP in HEK293 cells (210,000 copies), ORPs 10 and 11 are present at approximately 4,600 and 130,000 – roughly equal to their obligate binding partner, ORP9 at 130,000 copies (Cho et al., 2022). Therefore, assuming similar turnover numbers for ORPs9-11 and OSBP, they will be poised to turnover similar magnitudes of PI4P. Indeed, although experiments have shown that loss of ORP9/10 complexes leads to elevation of Golgi PI4P and enhanced recruitment of OSBP, the converse might equally be true: loss of OSBP may, in elevating PI4P, lead to enhanced recruitment and activity of ORP9/10! In other words, Golgi PI4P homeostasis in growing cells is likely maintained by the combined activities of OSBP and ORP9/10 or ORP9/11 complexes.

A quantitative consideration of Golgi PI4P turnover is not possible from our data, since we would need a robust calibration of membrane recruitment of our PI4P probes at defined PI4P mol fractions. However, we do know the turnover number of OSBP is approximately 0.5/s (Mesmin et al., 2013) and that there are approximately 210,000 copies of the protein (Cho et al., 2022), heavily enriched at the Golgi (**Fig. 5**). This places the turnover capacity of OSBP in a HEK cell at ∼100,000 PI4P molecules per second. The total number of PI4P and PI(4,5)P2 molecules in a cell is approximately equal, estimated to be around 10 million (Wills and Hammond, 2022), with perhaps half of this PI4P concentrated in Golgi membranes. Therefore, OSBP could be consuming as much as ∼2% of Golgi PI4P each second. However, the time to deplete PI4P after blocking synthesis will be complex, since OSBP requires PI4P for Golgi targeting, causing its turnover number to decline as it depletes PI4P. A similar argument with apply to ORP9/10/11 complexes. Nevertheless, at steady state OSBP would be able to turnover the entirety of the Golgi PI4P fraction in around a minute in a HEK cell. Since OSBP and ORPs 9-11 are ubiquitously expressed in all tissues, (Lehto et al., 2001) both likely contribute to similarly rapid PI4P turnover in all mammalian cell types, depending on their relative expression.

The essential role for OSBP in turnover of ectopic PI4P induced at mitochondria implies a wider role for OSBP in PI4P homeostasis than just Golgi pools, particularly since PI4P accumulation seems sufficient to trigger this (**Fig. 4**). It is for this reason that we tested for a role of OSBP in PI4P homeostasis in the membrane containing the most PI4P: the plasma membrane. However, we found no impact of OSBP on plasma membrane PI4P turnover, despite its localization there (**Fig. 6**). The most likely explanation is the extremely low density of particles, approximately 1 per 5 µm2 of plasma membrane. Contrast this with an estimated 100,000 molecules of PI4P in a similar membrane area (Wills and Hammond, 2022) and the estimated turnover number of OSBP (and other ORPs) at ∼0.5/s (Mesmin et al., 2013; Lipp et al., 2019). There is simply not enough OSBP present in the PM at any given moment to make a significant impact on PM PI4P turnover. The PI4P concentration in the plasma membrane is likely not sufficient to drive recruitment of enough OSBP without its co-receptor, Arf1 (Levine and Munro, 2002). We speculate that OSBP’s low density visits to ER/PM contact sites are not functional, but rather an accident of the high density of PI4P molecules in this compartment, the presence of VAP proteins and the large number of OSBP molecules.

In conclusion, our results demonstrate a significant impact of OSBP on Golgi PI4P levels. Therefore, not only does OSBP facilitate cholesterol traffic to the Golgi through anti-port with PI4P, it also contributes to Golgi PI4P homeostasis. It is thus likely an important regulator of non-ORP controlled functions in this organelle, such as vesicular traffic.

## Materials and Methods

### Cell Culture and transfection

HEK293A cells (ThermoFisher R70507) were maintained in DMEM (5.56 mM glucose, glutaMAX supplement, Thermo Fisher 10567022) supplemented with 10% fetal bovine serum (ThermoFisher 10438-034), 10 u/ml penicillin and 10 µg/ml streptomycin (Thermofisher 15140122) and 0.1% (v/v) chemically defined lipid supplement (ThermoFisher 11905031, included to supplement the sterols, polyunsaturated fatty acids and fat-soluble vatiamins that may become depleted from serum). T75 flasks were passaged twice weekly by rinsing once in sterile phosphate buffered saline and dissociation in 1 ml of TrpLE (ThermoFisher 12604039) before diluting 1:5 into fresh media. For imaging, cells were seeded at 25-50% confluence on 35 mm #1.5 glass bottom dishes (CellVis D35-20-1.5-N) pre-coated with 10 µg/ml entactin-collagen-laminin (Sigma 08-110) in DMEM. For siRNA transfection, 50 pmol ON-TARGETplus Human OSBP1 siRNA Smartpool (Horizon Discovery L-009747-00-0005) or ON-TARGETplus Non-targeting Control Pool (Horizon Discovery D-001810-10-05) were pre-complexed with 6.6 µl DharmaFECT1 (Horizon Discovery T-2001-01) in 400 µl serum-free medium for 20 minutes before adding to dishes. After 48 hours, cells were re-seeded into fresh dishes for cDNA transfection to reduce cell density to ∼50%. cDNA transfection was accomplished by mixing 1 µg of DNA pre-complexed with 3 µg lipofectamine 2000 (ThermoFisher 51985091) in 200 µl Opti-MEM (ThermoFisher 11668019) for 20 min before adding to the dish. After 4 h, transfection complexes were removed and replaced with 2 ml fresh medium.

### Gene editing

NG2-OSBP cells were generated as described by the OpenCell project (Cho et al., 2022). Briefly, an existing HEK293A cell line expressing the NeonGreen21-10 split fluorescent protein (Wills et al., 2023) was suspended in electroporation buffer R (Neon Transfection system, ThermoFisher) at 40,000 cells/µl. 5 µl of this suspension was mixed with 5 µl buffer R containing 100 pmol homology-directed repair template (synthesized as a ssDNA oligo, gcgacggcggctctcgtaggcggttccggtcttgtatctccaggcggcggcggctcatggcggcgacggaACCGAGCTCA ACTTCAAGGAGTGGCAAAAGGCCTTTACCGATATGATGGGTGGCGGCgctgagaggagtggtggggccaggcccggcagc cattgcagcacttggcggcggcggcgccggtccccca, IDT), 10 pmol gRNA (atggcggcgacggagctgag, ThermoFisher) and 10 pmol TruCut Cas9V2 (ThermoFisher, A36498). The mixture was electroporated using the Neon electroporation system (ThermoFisher, MPK5000) with a single 20 ms pulse at 1500 V, according to the manufacturer’s instructions. Thereafter, cells were recovered in 2 ml antibiotic-free growth medium in a 6-well plate, and then sorted by fluorescence activated cell sorting to isolate neon-green positive cells.

### Immunofluorescence

HEK293A or NG2-OSBP cells were seeded at approximately 50% confluence on 20 µg/ml entactin-collagen-laminin coated 8-well glass slides (MP biomedicals ICN6040805) and returned to the incubator. After one hour, they were fixed adding formaldehye (Fisher Scientific 50-980-487) to the medium to reach 4% final concentration (concentrated formaldehyde prepared in water and 10x PBS to make an isotonic solution). After 15 minutes at room temperature, wells were rinse 3 times with 50 mM NH4Cl in PBS to quench unreacted aldehyde, then permeabilized with 0.2% triton X-100 in PBS for 5 minutes. Next, they were rinsed three times in PBS and blocked for 30 minutes in blocking solution (PBS + 5% by volume normal goat serum, ThermoFisher PCN5000). Staining was for 1 hour with 1:500 anti-OSBP (Atlas Antibodies HPA039227) followed by 30 min with 1:200 AlexaFluor647 goat anti-rabbit secondary (ThermoFisher A-21245) in blocking solution. Cells were rinsed 4 times in PBS, once in miliQ water and mounted in ProLong Diamond (ThermoFisher P36961) and #1.5 coverglass. Images were acquired by confocal microscopy.

### Microscopy

For live cell imaging, media was replaced with 1.6 ml Complete HEPES-buffered Imaging Medium (CHIM) consisting of 10% fetal bovine serum, 1:1000 chemically defined lipid supplement and 25 mM NaHEPES, pH 7.4 in FLuoroBrite DMEM (ThermoFisher A1896702). Cells were imaged on a Nikon TiE microscope stand through a 1.45 NA plan-apochromatic 100x oil-immersion objective lens (Nikon). For confocal, the TiE was mounted with an A1R-HD confocal laser scanning microscope and imaged in resonant mode. Co-excitation of green (EGFP) and far-red (iRFP713 or Alexa647) fluorescence was achieved using the 488 and 640 nm laser lines of a fiber-coupled LUN-V combiner. To prevent cross talk, red (mCherry) fluorescence was excited separately using the 561 nm laser line in a subsequent line scan. Emitted light was collected through filters specific to blue (425-475 nm), green (505-550 nm), yellow-orange (570-620 nm), and far-red (650-850 nm) wavelengths. Confocal planes were obtained with a 1.2 Airy Unit pinhole (calibrated for the 640 nm channel), with 8 or 16 frame averaging.

For TIRFM, a Nikon motorized TIRF illuminator was used. Laser excitation was achieved through a fiber coupled four-line combiner (405, 488, 561, and 638 nm) from Oxxius. Emitted light was collected using dual-pass filters from Chroma, specifically for blue/yellow-orange (420-480 nm / 570-620 nm) and green/far-red (505-550 nm / 650-850 nm) wavelengths mounted in a motized filter wheel (Sutter). Images were collected with a Hammamatsu ORCAfusion BT camera; single molecule imaging employed the ultra-low noise mode with 2x2 binned pixels and the microscope’s 1.5x magnifier lens.

Time lapse imaging on either system was accomplished using a motorized stage to select 10-12 fields on the dish for repeated imaging at a frequency of 2 per minute. For drug additions, compounds diluted in 0.4 ml CHIM at 5-fold final concentration were added as indicated to cells in 1.6 ml media. These additions and their final concentrations were: 1 µM rapamycin (Fisher Scientific BP2963-1), 30 nM GSK-A1 (Sigma SML2453), 20 nM OSW-1 (Cayman Chemical 30310), rCD1 (SiChem SC-7000) and/or 1 µM FK506 (SiChem FK-506). Single molecule imaging was accomplished on single cells with a cropped camera sensor (256x256 pixels) to enable rapid imaging at 20 Hz.

### Quantitative Image Analysis

All analysis was accomplished using the open source image analysis software Fiji (Schindelin et al., 2012). Quantification of P4Mx1 fluorescence in different membrane compartments was performed by background subtracting the images using the mode pixel intensity (corresponding to the background signal outside of the cells) and defining regions of interest (ROI) for each cell. The mean pixel intensity was then measured within a binary mask of the selected compartment, normalized to the mean pixel intensity outside of that mask for each cellular ROI. Masks were generated using an automated thresholding algorithm on images of a marker for that target organelle (such as the Fis1-tail targeted to mitochondria), employing wavelet decomposition as described in detail previously (Wills et al., 2021). Alternatively, when no marker construct was employed (for the experiments reported in figures 1 and 2), we used manual thresholding of P4Mx1 fluorescence at the first frame to define Golgi-associated fluorescence and generate a binary mask of that threshold for each frame. This worked robustly for P4M fluorescence wherein the Golgi signal is much brighter than the signal associated with plasma membrane or endosomes and cytosol. For quantification of P4Mx2 in TIRFM images, intensity of the cellular footprint was simply measured at each time point, and normalized to the pre-stimulus mean.

For single particle analysis, we used the ThunderStorm plugin (Ovesný et al., 2014) for Fiji to detect single particles and quantify intensity and size statistics. For particle detection, we used the default detection settings calibrated for our specific camera’s A/D conversion rate, noise floor and pixel size. For analysis of particle enrichment at ER, clathrin-labelled structures or MAPPER-labelled contact sites, we used a custom Fiji macro. This first used ThunderSTORM to count particles in a 100 µm2 region of the cell’s footprint. It then generated a wavelet-derived binary mask of the marker image (as for confocal analysis described in the preceding paragraph), and used this to filter localizations falling inside the mask, compared to those falling outside. The fold enrichment was calculated as the ratio of number of localizations inside the mask to outside.

All data were plotted using Prism 9 (Graphpad), which was also used for the indicated statistical tests.

### Plasmids

Plasmids were obtained as cited, or else generated by polymerase chain reaction or gBlock synthesis (IDT) and assembled using HiFi assembly (New England Biolabs) according to the manufacturer’s instructions as shown in **Table 1**. siRNA resistant OSBP was generated by designing the OSBP ORF with silent mutants conferring resistance to the siRNA smart pool used.

**Table 1:** Plasmids used in this study.

| Plasmid | Backbone | Insert | Reference |
| --- | --- | --- | --- |
| NES-EGFP-P4Mx1 | pEGFP-C1 | *X. laevis* map2k1.L(32-44):EGFP:*L. pneumophila* SidM<sup>546-647</sup> | (Zewe et al., 2018) |
| NES-iRFP-P4Mx1 | piRFP713-C1 | *X. laevis* map2k1.L(32-44):EGFP:*L. pneumophila* SidM<sup>546-647</sup> | (Zewe et al., 2018) |
| EGFP-P4Mx2 | pEGFP-C1 | EGFP:*L. pneumophila* SidM(546-647):SidM<sup>546-647</sup> | (Hammond et al., 2014) |
| mCherry-FKBP | pmCherry-C1 | mCherry:FKBP1A(3-108):[GGSA]4GG | (Hammond et al., 2014) |
| mCherry-FKBP-OSBP (siRNA resistant) | pmCherry-C1 | mCherry: SGLRSRSAAAGAGGAARAALG: FKBP<sup>3-108</sup>:SA(GGSA)<sub>4</sub>PRAQASNS:OSBP | This study |
| FRB-mito | piRFP713-C1 | iRFP: MTOR<sup>2021-2113</sup>: [GGSA]<sub>2</sub>QASNSAVDGTGTA: FIS1<sup>122-152</sup> | This study |
| SNAP-mito | piRFP713-C1 | iRFP: SNAP-tag®: [GGSA]<sub>2</sub>QASNSAVDGTGTA: FIS1<sup>122-152</sup> | This study |
| pAcGFP-Sec61β | piRFP-C1 | pAcGFP:SEC61B | Addgene plasmid #15108 (Voeltz et al., 2006) |
| EGFP-VAPB | pmCherry-C1 | mCherry:VAPB | (Zewe et al., 2018) |
| mCherry-GRIP | pmCherry-C1 | mCherry: SGLRSRAQASNSAVD:GOLGA4<sup>V2162-S2243</sup> | This study |
| mCherry-FKBP-PI4KA<sup>C1001</sup> | pmCherry-C1 | mCherry: TagBFP2: FKBP1A3-108:[GGSA]4GG:PI4KA<sup>1102-2103</sup> | (Zewe et al., 2020) |
| mCherry-FKBP-PI4KA<sup>C1001-D1957A</sup> | pmCherry-C1 | mCherry: TagBFP2: FKBP1A3-108:[GGSA]4GG:PI4KA<sup>1102-2103-D1957A</sup> | (Zewe et al., 2020) |
| mCherry-MAPPER | pmCherry-C1 | mCherry:MAPPER | (Chang et al., 2013) |
| mCherry-CLC | pmCherry-C1 | mCherry:CLTA | This study |

## Acknowledgements

We are grateful to Jen Liou (UT SouthWestern, USA) for sharing EGFP-MAPPER. We are grateful to all members of the Hammond lab for technical assistance with experiments during restrictive COVID mitigation. This works was supported by NIH grant R35GM119412. All authors declare no competing financial interests.

